# Isotope Dilution nanoLC-MS/MS Quantitation of Methylglyoxal DNA-Protein Crosslinks: Formation and Repair in Human Cells

**DOI:** 10.1101/2025.10.28.685209

**Authors:** Reinner O. Omondi, Elijah M. Barnes, Krishna C. Gurajala, Gabrielle Fisette, Ibaad A. Chaudray, Luke Erber

## Abstract

DNA-protein crosslinks (DPCs) represent a prevalent form of DNA damage that form when cellular proteins become covalently trapped to DNA strands upon exposure to various endogenous and exogenous agents. Methylglyoxal, is an endogenous metabolite that reacts with guanine and adenine bases in DNA and RNA, as well as cysteine, arginine and lysine residues in proteins, generating advanced glycation end-products (AGEs) including DPCs. These modifications have been linked to human disease, including cancer, liver disease, diabetes, and neurodegenerative disorders. Herein, we present a mass spectrometry method for quantifying MGO-induced DNA-protein crosslinks (DPCs) in human cells. We prepared an isotope ^15^N_2_^13^C_6_-dG-MGO-Lys internal standard to develop a quantitative LC-MS/MS method for detecting and quantifying the formation and repair of dG-MGO-Lys DPCs in cells. Genomic DNA was extracted, subjected to sequential protease and nuclease digestion, purified by offline HPLC, and analyzed by LC-MS/MS. The method’s standard curve showed a strong linear relationship across a concentration range of 10-1000 fmol (R^2^ = 0.9994). The method achieved limits of detection (LOD) and quantification (LOQ) of 10 and 20 fmol, respectively. Inhibition of proteasome and SPRTN activity revealed that SPRTN functions as a predominant proteolytic enzyme in MGO DPC repair. Overall, this analytical approach can offer valuable insights into the relevance of DPCs in diseases linked to elevated MGO levels.

## INTRODUCTION

The human genome is exposed to various forms of DNA damage. DNA-protein crosslinks (DPCs) are a common type of DNA damage and form when cellular proteins become covalently trapped to DNA strands upon exposure to various endogenous and exogenous agents.^1^ DPCs can be induced by exogenous agents, including ionizing radiation,^2^ UV light,^3^ transition metal ions,^4^ environment contaminants,^5^ and common anticancer agents such as platinum compounds,^6^ nitrogen mustards,^7^ and haloethylnitrosoureas.^8^ Endogenously, DPCs form upon exposure to reactive oxygen species, ^9, 10^ and reactive metabolites such as formaldehyde and methylglyoxal.^11-13^ Owing to their bulky structure, DPCs obstruct fundamental DNA metabolic processes such as DNA replication, repair, transcription, and recombination.^14, 15^ DPC formation has been associated with premature aging, cancer predisposition, cardiovascular disease, and neurodegenerative disorders.^13^

DPC repair is thought to proceed in two general steps. First, proteolytic enzymes such as SPRTN and the proteasome to generate DNA-peptide crosslinks or DNA adducts.^16, 17^ The importance of proteolysis in DPC repair pathway is highlighted by hypomorphic mutations in the SPRTN gene in individuals with Ruijs-Aalfs syndrome. Patients with Ruijs-Aalfs exhibit elevated levels of DPCs, increased genomic instability, premature aging, and are at higher risk for developing hepatocellular carcinoma.^18^ Other proteolytic factors implicated in DPC repair include FAM111A,^19^ DDI1, DDI2,^20^ and ACRC.^1, 21, 22^ The proteolyzed DPC products are subsequently processed by DNA repair mechanisms including nucleotide excision repair (NER),^23^ homologous recombination (HR),^24^ or translesion synthesis (TLS) polymerases.^25-27^ The importance of these DNA repair processes in DPC repair are highlighted by studies of loss-of-function mutations in XRCC8 and ERCC6 (encoded by CSA and CSB genes respectively) which participate in NER.^28^ Cells with loss of CSB and CSA activity exhibit reduced DPC repair capacity and this loss may contribute to unique pathological features of Cockayne Syndrome including cachexia, neurodegeneration and kidney failure. Alternatively, nucleases such as MREII,^29^ APEX1 and APEX2^30^ are observed to process DPCs in the absence of proteolysis. Additionally, post-translational modifications such as ubiquitination and SUMOylation regulate protease recruitment and processing efficiency.^31, 32^

Methylglyoxal (MGO) is a highly reactive α-oxoaldehyde capable of forming a wide range of cyclic and acyclic DNA adducts.^33^ Endogenously, MGO is generated as a by-product of glycolytic pathway, by degradation of triosephosphates, dihydroxyacetone phosphate and glyceraldehyde-3-phosphate or through nonenzymatic sugar fragmentation.^34^ MGO can also form via ketone body oxidation, lipid peroxidation and amino acid catabolism.^34^ Although MGO is present in various commonly consumed foods, exogenous intake is unlikely to contribute significantly to plasma MGO levels.^35^ Notably, its endogenous formation rate varies with organism, tissue type, cellular metabolism, and physiological state.^36^ Reported plasma concentrations range from approximately 100 nM to 400 μM.^37, 38^ MGO metabolism is achieved through the glyoxalase system, wherein lactoylglutathione lyase and hydroxyacylglutathione hydrolase (GLO1 and GLO2 respectively) convert MGO to D-lactic acid.^39, 40^ Due to its high reactivity, MGO readily modifies DNA bases and protein residues, leading to the formation of advanced glycation end-products (AGEs).^41^ DNA products include N^2^-carboxyethyl-2’-deoxyguanosine (CEdG), and 3-(2′-deoxyribosyl)-6,7-dihydro-6,7-dihydroxy-6/7-methylimidazo-[2,3-b]purine-9-one adducts (cMG-dG) (Fig. 1). Representative protein adducts include N*ε*-(1-carboxyethyl)-L-lysine (CEL) and N^6^-(N-(1-carboxyethyl)carbamimidoyl)-L-lysine (CEA) and the lysine dimer, 1,3-di(N*ε*-lysino)-4-methyl-imidazolium (MOLD) (Fig. 1). ^42^ In addition, MGO has also been shown to generate covalent DNA-protein crosslinks (Fig. 1), ^43, 44^ and protein-protein crosslinks.^45^ These modifications disrupt cellular homeostasis and are associated with numerous pathologies, including cancer,^46^ diabetes,^47^ obesity,^48^ and cardiovascular and renal diseases.^49, 50^ Recent work has demonstrated that identification and quantitation of MGO adducts serve as improved predictors for diabetic kidney disease.^51^

**Figure 1:**
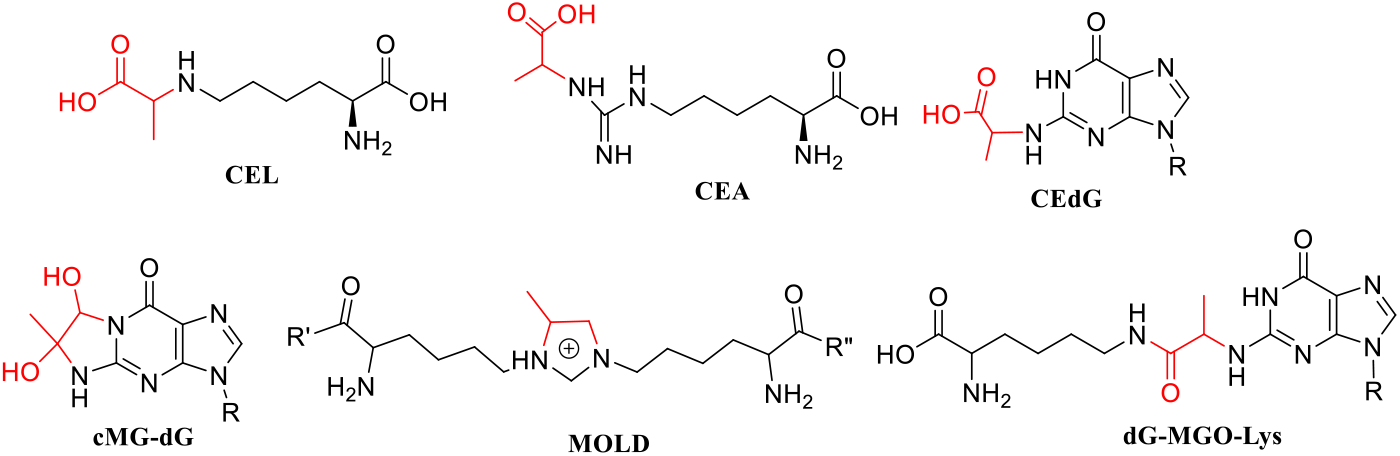
Chemical structures of MGO-modified nucleosides and lysine residues. dR represents the deoxyribose sugar, while R’ and R″ denote hydroxyl groups.

Our recent findings demonstrated that reduced expression of the SPRTN gene resulted in elevated DPC accumulation following MGO treatment, thereby supporting a role for SPRTN in the recognition and removal of MGO-derived DPCs.^12^ However, the direct quantification of specific MGO-induced DPCs in human cells remains limited. Previous studies had indicated that dG and lysine crosslinked in the presence of MGO to form dG-MGO-Lys, ^44, 52^ and we hypothesized that this crosslink serves as a specific DPC biomarker of MGO metabolism. Therefore, the present work aims to develop a sensitive liquid chromatography-tandem mass spectrometry (LC-MS/MS) assay to specifically measure the formation and repair of the dG-MGO-Lys conjugate by employing an isotopically labeled dG-MGO-Lys as an internal standard. Our method is robust, with a standard curve displaying strong linearity and LOD and LOQ of 10 and 20 fmol, respectively. In vitro experiments confirm that MGO-derived DPCs are persistent and are repaired through SPRTN proteolytic activity. Understanding the formation and repair of MGO-derived DPCs has the potential to provide a valuable biomarker for predicting disease risk, onset, and progression.

## EXPERIMENTAL SECTION

### dG-MGO-Lys Crosslink Adduct Extraction3

Cells cultured in 15-cm plates were treated with MGO in serum free media, washed with 5 mL cold PBS, and lysed in 1 mL DNAzol. Proteinase K digestion was carried out at 37 °C for 1 h, with intermittent mixing. Samples were then chilled on ice, and DNA was precipitated with cold ethanol, pelleted by centrifugation (14,000 x g, 5 min, 4 °C), washed twice with 70% ethanol, air-dried, and dissolved in 400 μL TE buffer for quantification using PicoGreen. 1 μg of DNA was digested with 40 mM ammonia acetate (pH 6.0) containing 10mM CaCl_2_, and 1.4 U/mL pronase (2 μL), with the reaction volume adjusted to 100 μL and incubated overnight at room temperature. The digested DNA was ethanol precipitated and resuspended in 100 μL digestion buffer containing 40 mM ammondia acetate (pH 6.0), 10mM CaCl_2_, 10 mM MgCl_2_ and an enzymatic cocktail including 42 mU phosphodiesterase I (0.6 μL/μg DNA), 24 mU phosphodiesterase II (0.4 μL/μg DNA), 330 mU alkaline phosphatase (0.2 μL/μg DNA), 7 mU DNAse (0.2 μL/μg DNA), 0.83 U/mL carboxypeptidase Y (5 μL), and 0.083 U/mL aminopeptidase M (5 μL). The reaction mixture was incubated overnight at 37 °C, spiked with 0.1 pmol isotope labeled MGO-DPC as an internal standard. Digests were filtered through Nanosep 10K devices, centrifuged, and transferred to a HPLC vial for downstream analysis.

### dG-MGO-Lys Crosslink HPLC Purification

dG-MGO-Lys was purified using an Agilent 1260 Infinity II HPLC system. The separation was carried out using buffer A (0.05% acetic acid in water) and buffer B (0.05% acetic acid in acetonitrile) on a Waters Atlantis T3 column (3 μm, 150 mm × 4.6 mm i.d.) at a flow rate of 0.4 mL/min. The gradient began at 2% B for 3 min, increased to 9% B at 21 min, ramped to 90% B at 23 min, and was held for 5 min before returning to 2% B at 30 min, followed by a 5-min equilibration. One-minute fractions were collected. The dG-MGO-Lys adduct fractions, A (22.70 - 23.70 min) and B (23.70 - 24.70 min) were dried under vacuum, resuspended in 0.1% formic acid, and sonicated. The samples were then transferred into a glass vial for subsequent LC-MS/MS analysis.

### DNA Quantification by UV-HPLC

To normalize crosslink levels between samples, DNA quantification was carried out by using an external a deoxyguanosine (dG) calibration curve. A 1.76 mM dG solution was prepared. Known mole amounts of dG were injected into the HPLC and processed according to the method described for the crosslink HPLC purification. The UV peak area observed at 20.96 min using 254 nm UV wavelength was recorded for dG quantification. dG levels from samples were calculated by fitting to a calibration curve constructed from the injection of known dG amounts.

### Quantitation of dG-MGO-Lys Adduct by LC-MS/MS

Quantitative analysis of dG-MGO-Lys was performed using a Vanquish Neo nanoHPLC system coupled with a Altis Plus triple Quadrupole mass spectrometer (Thermo Scientific). Liquid chromatography separation was performed using buffer A (0.1% formic acid in water), and buffer B (0.1% formic acid in 80% acetonitrile). Analytes were separated with a self-packed fused silica capillary column (40 cm length, 75 μm inner diameter, 5 μm Zorbax C18 (Agilent)) using a flow rate of 0.350 μL/min. The separation gradient started at 1% B for one min before increasing to 30% B at min 15. The %B was raised to 90% B in 0.1 min. The gradient was held at 90% B for 4.9 min before dropping the %B to 1% at 21 min. The column was equilibrated for an additional 5 min at 1% B. The authentic standard for dG-MGO-Lys eluted at 11.46 and 11.74 mins. The mass spectrometry source conditions were set at 2.6 kV, capillary temperature of 275 °C. Mass spectrometry data was collected in positive polarity with a Q1 resolution of 0.7 FWHM, Q3 resolution of 0.2 FWHM, CID gas pressure set at 0.5 mTorr and a dwell time of 100 ms. Analysis of dG-MG-Lys was performed using multiple reaction monitoring (MRM) mode. MRM analysis was conducted by monitoring the characteristic fragmentations of the analyte, including the neutral loss of deoxyribose, protonated lysine ion, and the combined loss of the deoxyribose and lysine. The monitored transitions were *m/z* 468.2 [M + H]^+^→ 352.2 [M -116 + H]^+^, *m/z* 468.2 [M + H]^+^ → 206.1 [Lys +H]^+^, *m/z* 468.2 [M + H]^+^ → 147.1 [M - 116 - Lys +H]^+^. Corresponding mass transitions were also monitored for the ^15^N_2_ ^13^C_6_ - labeled internal standard, *m/z* 476.2 [M + H]^+^ → 360.2 [M -116 + H]^+^, *m/z* 476.2 [M + H]^+^ → 206.1 [Lys +H]^+^, *m/z* 476.2 [M + H]^+^ → 155.1 [M - 116 – Lys + H]^+^.

### Statistical Analysis

Statistics was conducted using Graphpad PRISM (version 10.1.0). Comparisons were assessed using ordinary one-way ANOVA using Turkey’s multiple comparisons test, with a single pooled variance. Group datasets statistics performed using Ordinary two-way ANOVA using uncorrected Fisher’s LSD, with a single pooled variance. GP: 0.1234 (ns), 0.0332 (^*^), 0.0021 (^**^), 0.0002 (^***^), <0.0001 (^****^).

## RESULTS AND DISCUSSION

### Synthesis and Characterization of ^15^N_2_^13^C_6_-dG-MGO-Lys Standard

We prepared an internal standard to establish an isotope dilution LC-MS/MS method to selectively and accurately measure dG-MGO-Lys DPCs in human cells. In this method the nucleoside-amino acid crosslinks are enzymatically released from the genome and spiked with a known amount of isotopically labeled standard. DPCs were purified by HPLC and analyzed using LC-MS/MS. Mass spectrometry analysis permits quantitation of the DPC as a light-to-heavy isotope ratio. To conduct this method, a heavy isotope labeled standard was synthesized following reported methods.^44^ Briefly, equimolar amounts of dG, MGO, and boc-protected ^15^N_2_^13^C_6_-L-Lysine were reacted under physiologically relevant conditions to yield the protected conjugate, which was subsequently subjected to catalytic hydrogenation over Pd/C to remove the protecting group (Fig. 2a). The compound was HPLC-purified and characterized by high resolution mass spectrometry. The total ion chromatogram displays two distinct peaks at 12.51 and 12.65 mins, indicating the formation of two isomeric species (Fig. 2b). Both full-scan and product-ion scan spectra exhibited the expected fragmentation patterns, confirming the formation of the conjugate, with a mass accuracy of <0.4 ppm.

**Figure 2:**
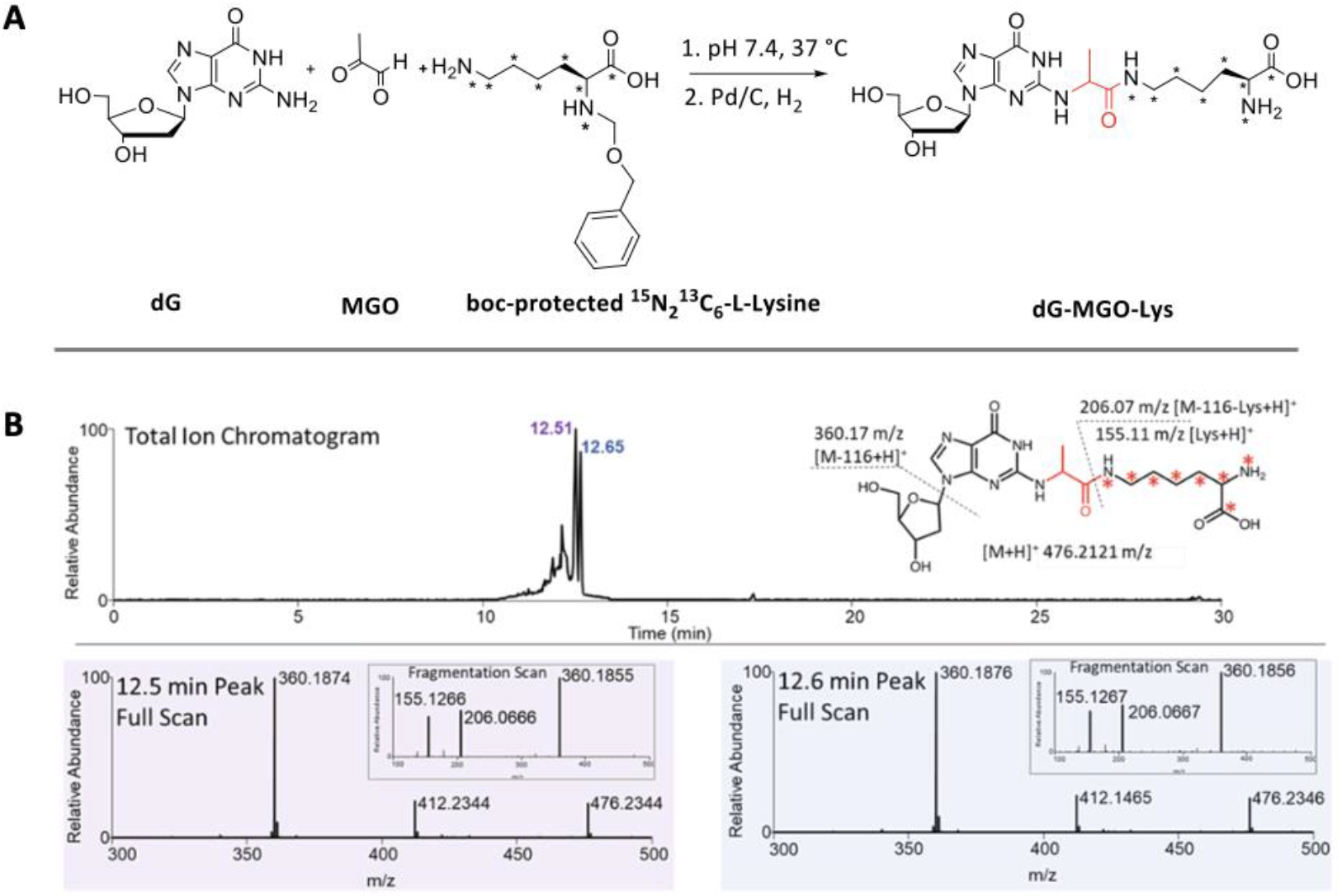
Synthesis and characterization of isotope ^15^N_2_ ^13^C_6_ - dG-MGO-Lys internal standard. **(a)** Synthetic route scheme. **(b)** LC-MS/MS analysis including ion chromatography and high-resolution MS/MS for peaks at 12.5 min and 12.6 min.

### LC-MS/MS Method Development for dG-MGO-Lys

We adopted a LC-MS/MS method to measure dG-MGO-Lys. This approach has been successfully used to quantify DPCs formed by formaldehyde, ^53^ and reactive oxygen species.^54^ When examining the ion chromatography, both the standard (dG-MGO-Lys) and the internal standard (IS) were observed at the same retention time (Fig. S1) We monitored three sets of transitions to improve the accuracy of our method. These transitions include the neutral loss of deoxyribose [M – 116 + H]^+^, the protonated lysine [Lys + H]^+^, and the combined loss of deoxyribose and lysine [M – 116 – Lys +H]^+^. For pure standards, the method achieved a limit of detection (LOD) was 5 fmol, and a limit of quantitation (LOQ) of 10 fmol (Fig. S2). The calibration demonstrated excellent linearity across three orders of magnitude (1-1000 fmol), with a R^2^ value of 0.9983.

### Quantification of dG-MGO-Lys DPCs in Cells

To identify and quantify dG-MGO-Lys from cells, we extracted genomic DNA from HeLa cells treated with 5 mM MGO for 2 h. We digested 1 µg DNA with proteases including proteinase K, pronase, carboxypeptidase Y, aminopeptidase M, and nucleases including DNase 1, alkaline phosphatase, phosphodiesterase I/II. The enzymatic digest was spiked with 100 fmol labeled internal standard, followed by HPLC purification and LC-MS/MS analysis of the HPLC purified samples (Fig. 3a). We observed formation of dG-MGO-Lys adducts in treated samples compared to untreated control samples (Fig. 3b). Using our optimized analytical HPLC method, we successfully resolved two peaks and both fractions (A and B) proved reliable for quantifying DPC levels in cells (Fig. S3 and S4). Inclusion of a RNAse treatment step prior to nuclease digestion of samples did not affect the overall dG-MGO-Lys yield (Fig. S5).

**Figure 3:**
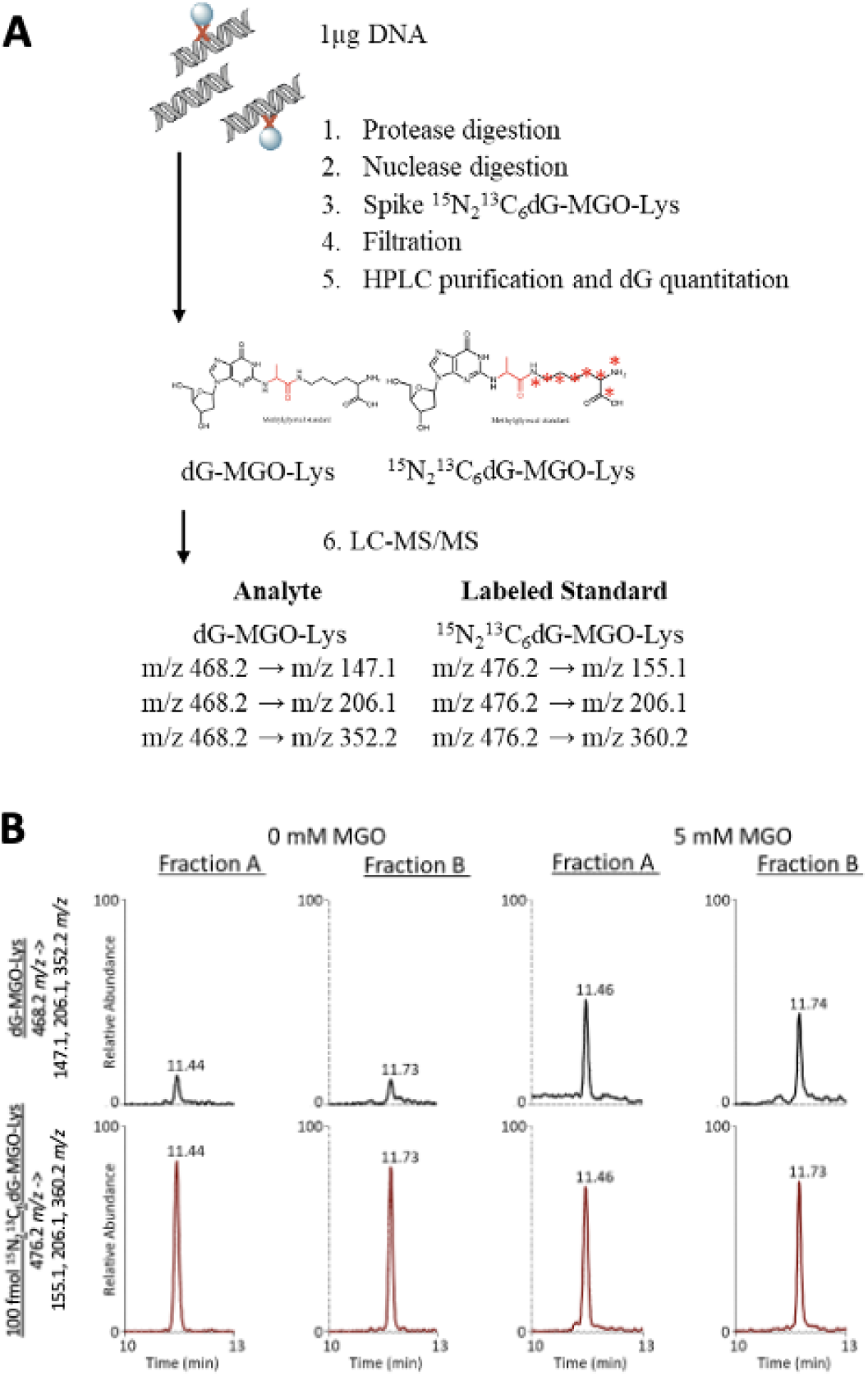
Extraction and measurement of dG-MGO-Lys crosslink adduct. **(a)** Experimental scheme for isotope dilution LC-MS/MS analysis of dG-MGO-Lys from genomic DNA **(b)** Representative ion chromatogram for HPLC fractions A and B for dG-MGO-Lys from untreated and MGO treated (5 mM, 2 h) HeLa cells.

### Calibration and Method Validation in DNA Matrix

To test the selectivity, accuracy, and sensitivity of our method, we developed a standard curve. Both the analyte and internal standard (IS) were detected at the same retention time. The lines of best fit for calibration standards were determined using linear least-square regression analysis based on the peak area ratios of the analytes to the IS. A strong linearity was observed in the concentration ranges from 10 to 1000 fmol, with an R^2^ values of 0.994 (Fig. S6). The accuracy of this method was calculated as 4, 11, and 11% for the high (50 fmol), middle (100 fmol) and low (500 fmol) validation points, respectively. The precision of the method was calculated as 5, 8 and 5% CV for the high, middle and low validation points, respectively (Table 1). Analysis of the spiked samples yielded LOD and LOQ at 10 and 20 fmol, respectively. Analyte recovery following filtration, HPLC purification and speedvac concentration was calculated by comparing the peak area of analyte spiked into control DNA spiked prior to sample processing with the peak area of same number of moles of IS spiked after speedvac concentration. The recovery was determined to be 34.4 ± 7.1%. We further assessed the chemical stability of the dG-MGO-Lys adduct in aqueous solution in an autosampler at 4 °C over a period of 72 h. The results showed that DPC levels remained unchanged. By comparing the peak area with and without the DNA matrix, signal suppression was quantified at 82.8 ± 14.7%.

**Table 1:** Method validation results for LC–MS/MS method for quantitation of dG-MGO-Lys in unexposed DNA (1 µg).

| moles (fmol) | Day | Mean | RSD (%) | Accuracy (%) | N |
| --- | --- | --- | --- | --- | --- |
| 50 (low) | 1 | $55.4 \pm 7.1$ | 13 | 11 | 3 |
| | 2 | $47.9 \pm 2.3$ | 5 | 4 | 3 |
| | 3 | $46.5 \pm 4.9$ | 11 | 7 | 3 |
| | Inter-day | $49.9 \pm 6.5$ | 13 | 0.2 | 9 |
| 100 (middle) | 1 | $90.7 \pm 8.2$ | 9 | 9 | 3 |
| | 2 | $89.0 \pm 7.0$ | 8 | 11 | 3 |
| | 3 | $91.6 \pm 0.8$ | 1 | 8 | 3 |
| | Inter-day | $90.4 \pm 6.3$ | 7 | 10 | 9 |
| 500 (high) | 1 | $406.1 \pm 13.6$ | 3 | 19 | 3 |
| | 2 | $414.9 \pm 10.3$ | 2 | 17 | 3 |
| | 3 | $428.9 \pm 13.7$ | 3 | 14 | 3 |
| | Inter-day | $416.6 \pm 15.7$ | 4 | 17 | 9 |

### Protease Digestion Efficiency

A challenge of this method lies in developing an enzymatic digestion protocol that fully releases the DPC adducts from the genome and produces small molecules suitable for MS/MS analysis while preserving the chemical identify of the DPC crosslinks. We set out to establish adequate digestion conditions by testing a series of proteases in different combinations (Fig. 4a). HeLa cells were treated with MGO (5 mM, 2 h) and DPCs were extracted according to the LC-MS/MS processing method. We compared five proteolytic digestion conditions including no protease, Proteinase K only, Pronase only, Carboxypeptidase Y and Aminopeptidase M only, or all four proteases together. Samples lacking protease treatment showed the lowest yield of dG-MGO-Lys. Notably, no appreciable difference was observed between proteinase K treatment and treatment with all four proteases, indicating that proteinase K alone was sufficient to achieve effective digestion.

**Figure 4:**
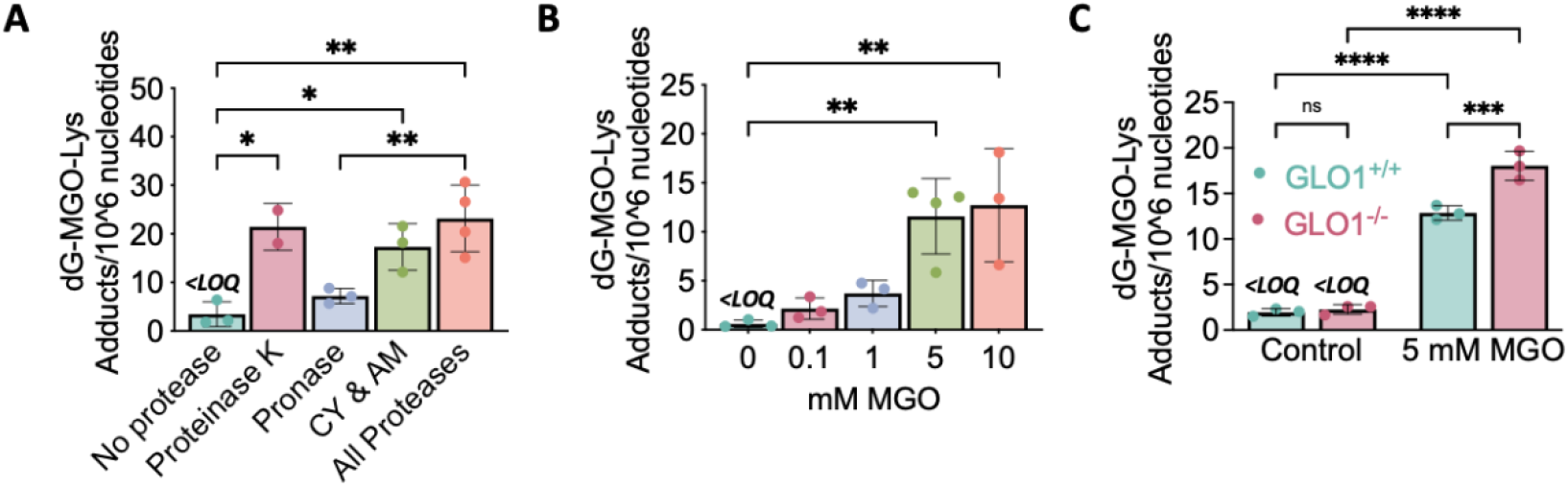
Quantitative analysis of dG-MGO-Lys DPC. **(a)**. HeLa cells treated with 5 mM MGO for 2 h were processed according to our LC-MS/MS method. Enzymatic processing was carried out as described in the method. Five proteolytic digestion conditions were compared, 1) no protease, 2) Proteinase K only, 3) Pronase only, 4) Carboxypeptidase Y (CY) and Aminopeptidase M (AM) only, 5) all four proteases together. **(b)** HeLa cells were exposed to increasing concentrations of MGO (0, 0.1, 1, 5 and 10 mM) for 2 h. dG-MGO-Lys DPC conjugates were quantified using the LC-MS/MS method. (c) Parental HEK293 cells or HEK293 cells deficient in GLO1 were treated with MGO (5 mM, 2 h). dG-MGO-Lys DPC conjugates were quantified using the LC-MS/MS method. Data are presented as mean ± SD (n = 3).

### Formation of dG-MGO-Lys DPCs

We applied the analytical method to examine the formation of MGO-induced DPCs under a range of MGO concentrations, including a more physiologically relevant concentration of 100 μM. Specifically, HeLa cells were treated with 0, 0.1, 1, 5 and 10 mM MGO for 2 h. We observed a pronounced dose-dependence increase in conjugate formation (Fig. 4b), from 2.16 ± 1.09 adducts per million nucleotides at 0.1 mM to 12.72 ± 5.7 adducts per million nucleotides at 10 mM. Treatment with 5 mM MGO yielded 11.59 ± 3.85 dG-MGO-Lys cross-links per million nucleotides, whereas DPC levels in the control group remained below LOQ. This is consistent with previously reported measurements of 2.1 dG-MGO-Lys adducts per million nucleotides in HT1080 cells MGO (5 mM, 2 h) exposure when quantified using an external curve.^12^ We hypothesized that inhibition of the glyoxalase system, specifically lactoylglutathione lyase (GLO1) would lead to elevated MGO DPC levels. We treated HEK293 parental and GLO1 knockout cells with MGO (5 mM, 2 h) and observed 12.87 ± 0.64 adducts per million nucleotides in parent cells and 18.05 ± 1.3 adducts per million nucleotides in GLO1 knockout cells.

### Repair of dG-MGO-Lys DPCs

The mechanisms by which cells recognize DPC lesions and direct them to specific repair pathways remain poorly understood. Given this type of DNA damage is uniquely formed by protein crosslinking, we examined the molecular basis of DPC proteolytic repair. We first established a controlled baseline by assessing the persistence of the dG-MGO-Lys adduct over 48 h after an acute MGO exposure using the ARK-SDS assay (Fig. 5a). The DPC levels remained 3.0 ± 1.0-fold higher than unexposed control at 36 h and 2.7 ± 0.1-fold higher at 48 h. To identify enzymatic pathways contributing to MGO DPC repair, we first evaluated the ubiquitin-proteasome pathway. In this pathway, ubiquitin molecules are covalently attached to proteins that are cross-linked to DNA, thereby tagging them as substrates for proteasomal processing.^56^ We treated cells with proteasome inhibitor MG132, E1 ubiquitin ligase UBA1 inhibitor TAK243, or a combination of both inhibitors (Fig. 5b). None of the treatments produced a significant increase in adduct levels compared to MGO treatment alone, suggesting that the ubiquitin-proteasome pathway is unlikely to represent a major repair mechanism for dG-MGO-Lys adducts. We also evaluated whether these DPCs could be recognized and processed by the SPRTN metalloprotease. To identify whether SPRTN is participating in repair of dG-MGO-Lys crosslinks, we generated a SPRTN knockout cell line (SK5) in HeLa cells (Fig. S7 and S8). The KO cell line was characterized by Sanger sequencing to reveal a five-nucleotide insertion resulting in a premature stop codon. TIDER analysis revealed half of the SPRTN gene copies were modified in this manner in this clone, resulting in a cell line exhibiting reduced SPRTN gene expression and proteolytic activity. We treated the SPRTN knockout and parental cells with MGO (5 mM, 2 h) and quantified the levels of dG-MGO-Lys DPC present (Fig. 5c). SPRTN-deficient cells exhibited a pronounced 2.27 ± 0.51-fold increase in DPC accumulation. We next examined the accumulation of DPCs in SPRTN-proficient and SPRTN-deficient cells under different treatments, including MG132 and inhibitors of ubiquitin (UBB) and SUMOylation, using the ARK-SDS assay (Fig. 5d). Inhibition of the proteasome with MG132 did not lead to any increase in DPC levels in either cell type. In contrast, blocking ubiquitination or SUMOylation produced increased DPC accumulation in SPRTN-deficient cells, 1.94 ± 0.26-fold and 1.86 ± 0.07-fold respectively. These differences in repair response to inhibition of ubiquitin and SUMOylation activity highlights the critical role of these post-translational modifications in regulating DPC resolution. Collectively, these results indicate that SPRTN, ubiquitination, and SUMOylation function in concert to facilitate efficient clearance of MGO DPCs.

**Figure. 5:**
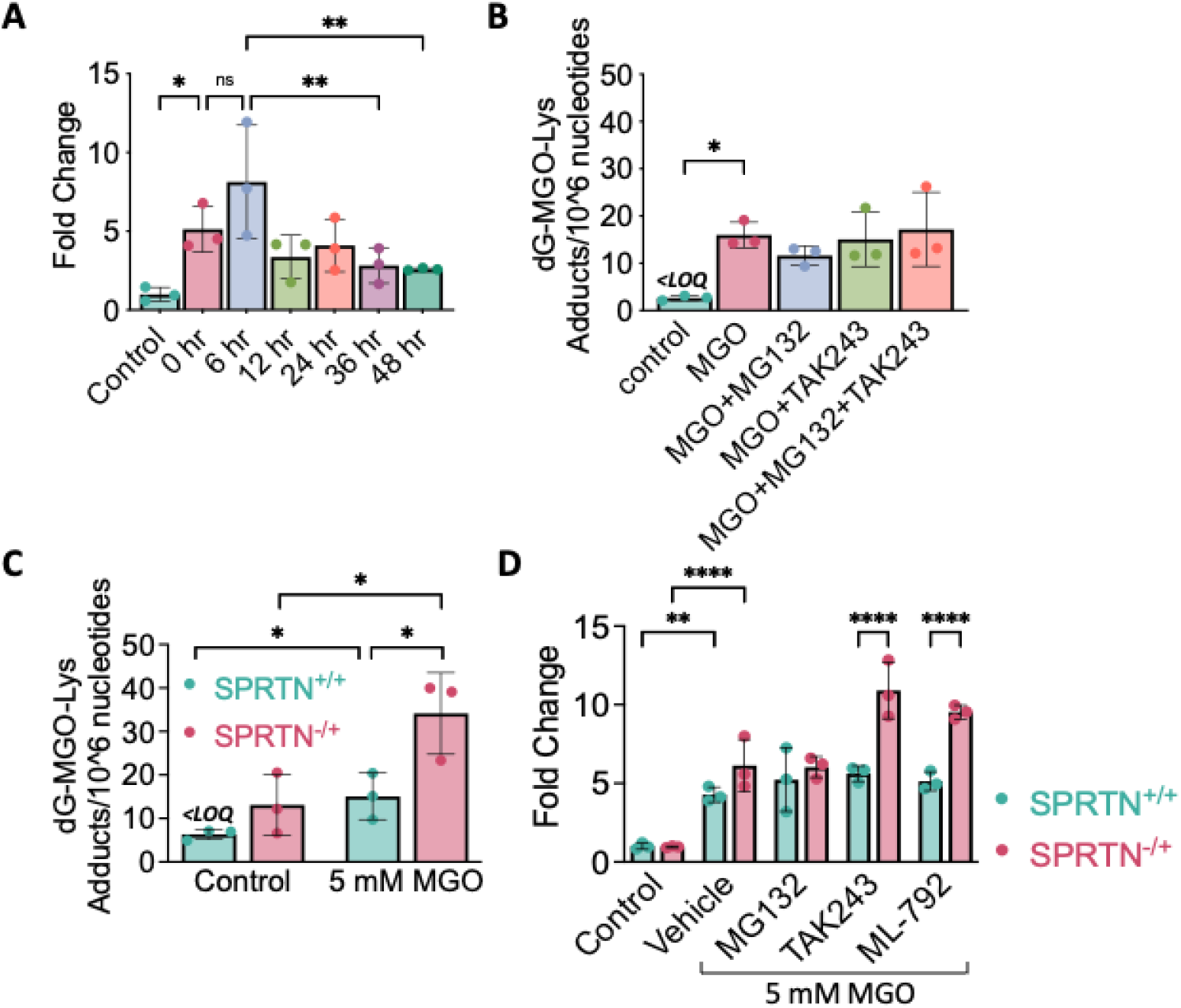
Repair of dG-MGO-Lys DPCs. **(a)** HeLa cells were treated with 5 mM MGO for 2 h. After removal of MGO-containing media and 2x PBS washes, cells were collected at initial, 6 h, 12 h, 24 h, 36 h, and 48 h post-treatment. DPCs were measured using ARK-SDS method. **(b)** HeLa cells were treated for 2 h with either vehicle, 5 mM MGO or in combination 1) 5 mM MGO and 10 µM MG312, 2) 5 mM MGO and 10 µM TAK243, 3) 5 mM MGO, 10 µM TAK243 and 10 µM MG312. dG-MGO-Lys was quantified from genomic DNA extracted from these treated samples. **(c)** SPRTN deficient and proficient HeLa cells were treated with 5 mM MGO for 2 h. dG-MGO-Lys was quantified from genomic DNA extracted from these treated samples. **(d)** SPRTN deficient and proficient HeLa cells were pretreated for 1 h with either vehicle, 1) 10 µM MG312, 2) 10 µM TAK243 or 3) 10 µM ML-792. Treated cells were then further treated with MGO in combination with the initial treatment for an additional 2 h. DPCs were measured using ARK-SDS method. Data are presented as mean ± SD (n = 3).

## DISCUSION

Accurate identification and quantification of DPCs are critical for understanding their biological impact and repair mechanisms. Several analytical methods have been developed to study DPCs. Classical biochemical approaches, such as KCl-SDS, ARK-SDS, modified comet assay, STAR, Western blotting and immunofluorescence, allow global assessment of cross-linking frequency and identification of the proteins and genomic regions participating in crosslinking. ^57-59^ However, these methods lack structural specificity.^57^ Mass spectrometry permits absolute quantitation of DPC levels in cells and tissues in a manner that offers high sensitivity, structural specificity and accurate quantification. Incorporation of stable isotope dilution strategies further enhance these measurements by correcting for matrix effects and sample processing losses.

Since DPC sample preparation and mass spectrometry-based processing of DPCs is laborious and requires specialized equipment, we sought to simplify the method where possible. We adopted the DPC extraction method described by the Swenberg lab to quantify DPCs formed by formaldehyde.^53^ Incomplete proteolytic digestion can lead to underestimation of adduct levels and is harder to monitor during sample processing. We anticipated incomplete digestion would leave cross-linked material intact, whereas overly aggressive conditions would risk nonspecific cleavage or degradation of analytes. We evaluated multiple proteolytic digestion approaches, including the use of proteinase K, pronase, aminopeptidase M, carboxypeptidase Y, and a combination of all proteases to optimize this step. We observed that utilizing a single proteinase K digestion (1 h at 37 °C) was sufficient. Compared to the method described by Swenberg’s lab to quantify the crosslink formed by formaldehyde, we added phosphodiesterase II and removed prolidase from mixture to ensure complete nuclease digestion. Important checks during sample processing included DNA quantitation by the PicoGreen kit to monitor genomic DNA yield from cell lysates. We monitored dG retention time and levels within our samples to monitor HPLC performance and to normalize DPC quantitation based on actual sample DNA µg amounts. To ensure effective nuclease digestion, we did not process samples exhibiting hydrophobic nucleotide oligomers, instead optimal digests exhibited only nucleoside products by HPLC-UV. Thus, establishing complete and reproducible digestion conditions is essential to ensure that LC-MS/MS measurements of DPCs accurately reflect their true abundance in biological samples.

In our efforts to characterize and quantify MGO-induced DPCs, we performed method validation to benchmark the LC-MS/MS analysis approach for the dG-MGO-Lys adduct. This method exhibited a LOD of 10 and LOQ of 20 fmol, values that are higher than reported for other DPC adducts. Swenberg and the group reported a lower LOQ of 37.5 amol using LC–MS/MS for the measurement of formaldehyde-induced DPCs, specifically, dG-Me-Cys adducts.^60^ The Tretyakova group reported a LOD of 2 fmol on column for reactive oxygen species-induced Thymine-Tyrosine DPC adduct.^61^ Given the LOQ for analysis of the dG-MGO-Lys standards in the absence of matrix is 10 fmol, we suspect that the differences in LOQ across the DNA adducts may be due to differences in ionization efficiency or chromatography. Future effort could be employed to optimize the chromatography conditions to improve the sensitivity of the dG-MGO-Lys LC-MS/MS method. Furthermore, we compared the levels of dG-MGO-Lys with DPC adducts observed in other cell culture-based studies. When human fibrosarcoma HT1080 cells were treated with mechloethamine (100 μM, 3 hours) or nornitrogen mustard (500 μM, 3 hours), 0.8 Cys-N7G-EMA adducts per million nucleotides and 0.13 Cys-NOR-N7G adducts per million nucleotides respectively were observed. ^62, 63^ We observe MGO DPC adducts form at a higher frequency of 2.16 adducts per million nucleotides when HeLa cells were exposed to 0.1 mM MGO for 2 h. These findings indicate a high frequency of DPC adducts in cells. Roughly a dozen protein and DNA adduct formed by MGO activity have been described. These adducts vary in tissue distribution, stability and LC-MS/MS measurements are complicated by limited availability of stable isotope standards. One of the major DNA adducts formed when MGO reacts with double stranded DNA is CEdG. This reaction introduces a chiral center and CEdG is observed as a diasteromeric mixture that is readily separated by HPLC and LC-MS/MS.^64^ Both isomers were observed in human breast tumor and adjacent normal tissue samples. In this study we observed baseline separation of two isomeric peaks of dG-MGO-Lys when processing DNA from cell samples. These diasteromers were consistent with the diastereomeric products formed when we prepared the isotope labeled standard. Notably, both peaks could be reliably employed for quantification, significantly improving the accuracy of the method. Future observations of the diastereomer ratio in vivo may reveal stereochemical biases in DPC repair. We compared the levels of dG-MGO-Lys with other MGO-derived adducts. Adduct levels will depend on factors including cell type, tissue, species, and dosage of exposure. When human acute promyelocytic leukemia (HL-60) cells were exposed to 524 µM MGO for 2 h, 18.2 ± 6.5 cMG-dG adducts per million nucleotides and 1.33 ± 1.12 CEdG adducts per million nucleotides were measured.^65^ The most similar MGO exposure level (1 mM, 2 h) in HeLa cells resulted in 3.1 ± 1.4 DPC adducts per million nucleotides. These findings place dG-MGO-Lys adduct at a similar level as the stable CEdG adduct. Finally, molecular studies of methylglyoxal metabolism have place GLO1^66^ among the enzymatic contributors to methylglyoxal abundance within cells. Our findings show that inhibition of this pathway also have the potential to contribute to DPC formation as a possible mechanism of MGO toxicity.

Development of DPC-selective quantitative approaches is important in dissecting the DNA repair response upon exposure to elevated levels of endogenous reactive molecules such as formaldehyde, methylglyoxal and reactive oxygen species. We recently used LC-MS/MS to demonstrate genomic DNA from tissues of SPRTN deficient mice exhibited elevated ROS-induced thymidine-tyrosine DPC levels in kidney, liver, heart and brain.^13^ In our initial study of MGO-induced DPCs, we observed that reduced expression of the SPRTN gene resulted in elevated DPC levels following treatment with 2.5 and 5 mM MGO.^12^ Consistent with these results, our findings using LC-MS/MS further support the notion that SPRTN may serve as a predominant repair enzyme responsible for resolving these lesions. The results from both LC-MS/MS and ARK-SDS assays indicate that the proteasome is likely to be a minor contributor for repair of dG-MGO-Lys adducts. A previous study reported that MGO can directly disrupt the proteasome through two routes. MGO modifies the 20S proteasome subunits and reduces their catalytic activity as well as downregulates 19S regulatory proteins, which compromises substrate recognition and processing.^68^ In our current study, inhibition of E1 ubiquitin ligase UBA1 or SUMO ligase SAE in SPRTN-deficient cells led to increased accumulation of DPCs. This observation is consistent with an earlier dot blot analysis, which showed a dose-dependent increase in ubiquitin and SUMOylation signals in MGO-derived DPCs.^12^ In the earlier study, complementary mass spectrometry–based proteomics also identified ubiquitin isoforms and SUMO proteins among the species covalently trapped on DNA, reinforcing their likely involvement in the processing of these lesions. Since our SPRTN knockout is only partial, we are unable to fully determine whether the ubiquitination and SUMOylation also contributes to regulation of other DPC repair mechanisms. Future investigations could include a SPRTN inducible knockout cell model or other proteases including FAM111A, DDI1 and DDI2, and ACRC.

## CONCLUSION

We developed and validated a isotope dilution nanoLC-MS/MS method to identify and quantify MGO-mediated DPCs in human cells. Using this approach, we demonstrated that DPC abundance is modulated by both metabolic and proteolytic activities. Among the proteolytic pathways examined, SPRTN appears to be a predominant proteolytic enzyme responsible for repairing MGO-induced DPCs, demonstrating its critical role in maintaining genomic stability. Our observations also support an important role for ubiquitination and SUMOylation in DPC repair. Beyond offering a sensitive and selective quantification strategy, this LC-MS/MS method has the potential to provide valuable insights into DPCs as potential biomarkers of MGO-associated pathologies, such as cancer and neurodegeneration.

## AKNOWLEDGEMENTS

We thank members of the Erber lab and Alex Hurben for their insightful suggestions and discussion. Support for this work was provided by startup funds from the University of Kansas (PI: Erber, L) and funded by a grant from NIGMS of the National Institutes of Health (R35GM159935, PI: Erber, L). Support for synthesis of the isotope labeled standard was provided through the National Institutes of Health grant (P30GM145499) to the University of Kansas’ Center for Molecular Analysis of Disease Pathways (CMADP), an NIH Center of Biomedical Research Excellence (COBRE).

## Notes

The authors declare no competing financial conflict

