## Supplemental Figures for "Isotope Dilution nanoLC-MS/MS Quantitation of Methylglyoxal DNA-Protein Crosslinks: Formation and Repair in Human Cells"

### Table of Contents


| | | |
|---|---|---|
| I. | Supplementary Figures | Page 3 |
| II. | Materials and Methods | Page 10 |
| III | References | Page 12 |

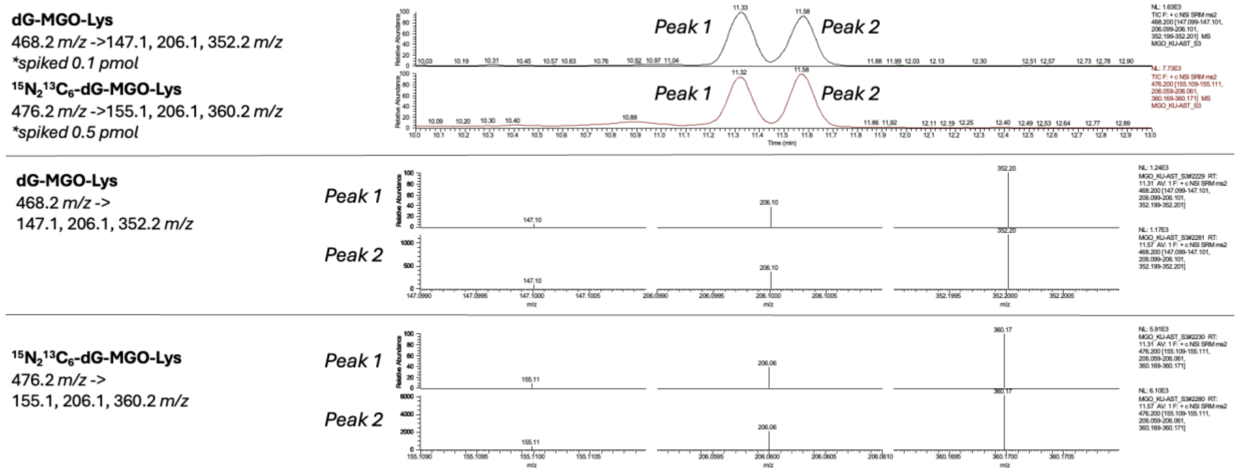


**Figure S1.** Structural verification of dG-MGO-Lys standards. (**a**) Representative LC-MS/MS ion chromatography for dG-MGO-Lys unlabeled and isotope labeled standards. (**b**) Representative $MS^2$ fragmentation pattern of dG-MGO-Lys standards from precursor ions represented in the ion chromatography.

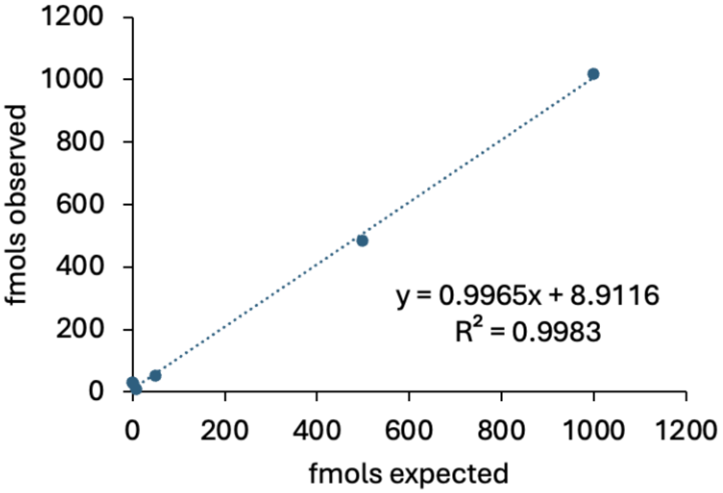


**Figure S2**. Calibration curve for the authentic dG-MGO-Lys standards spiked in aqueous buffer (0.1% formic acid in water). Observed signal (fmol) of unlabeled standard plotted against the signal for the isotope labeled standard across concentration range of 1-1000 fmol. Linearity, Limit of Detection (LOD), and Limit of Quantitation (LOQ) were calculated from these observations.

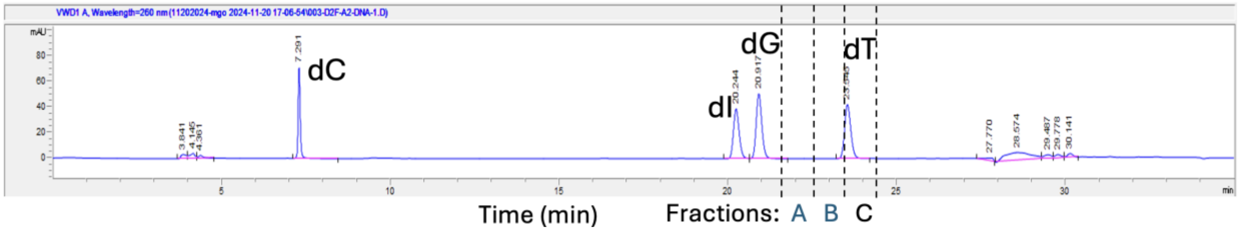


**Figure S3.** HPLC chromatogram of nucleoside separation. Representative separation of deoxycytidine (dC) at 7.2 min, deoxyinosine (dI) at 20.2 min, deoxyguanosine (dG) at 20.9 min, and thymidine (dT) at 23.5 min using UV detection at 260 nm. Fractions A, B, and C (indicated by dashed lines) correspond to the collected retention windows for subsequent LC-MS/MS analysis.

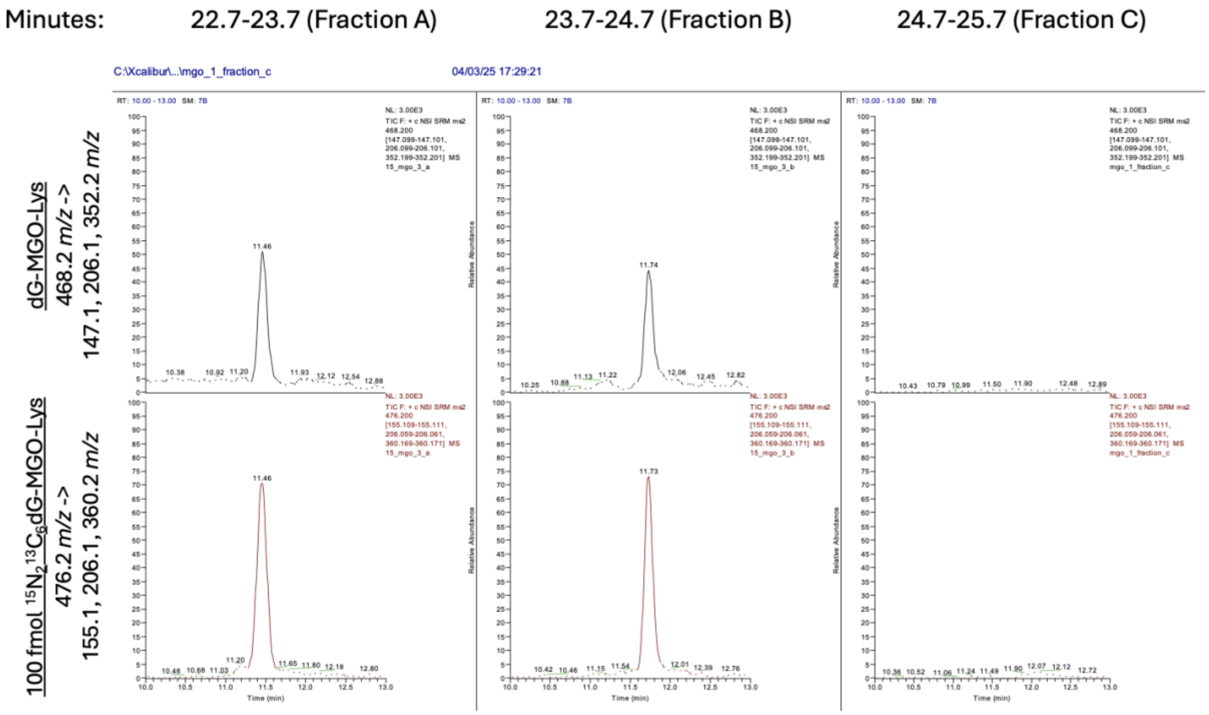


**Figure S4**. LC-MS/MS analysis of collected HPLC fractions. Representative extracted ion chromatograms of unlabeled dG-MGO-Lys and isotope $^{15}N_2{}^{13}C_6$- dG-MGO-Lys standard from HPLC fractions A (22.7–23.7 min), B (23.7–24.7 min), and C (24.7–25.7 min).

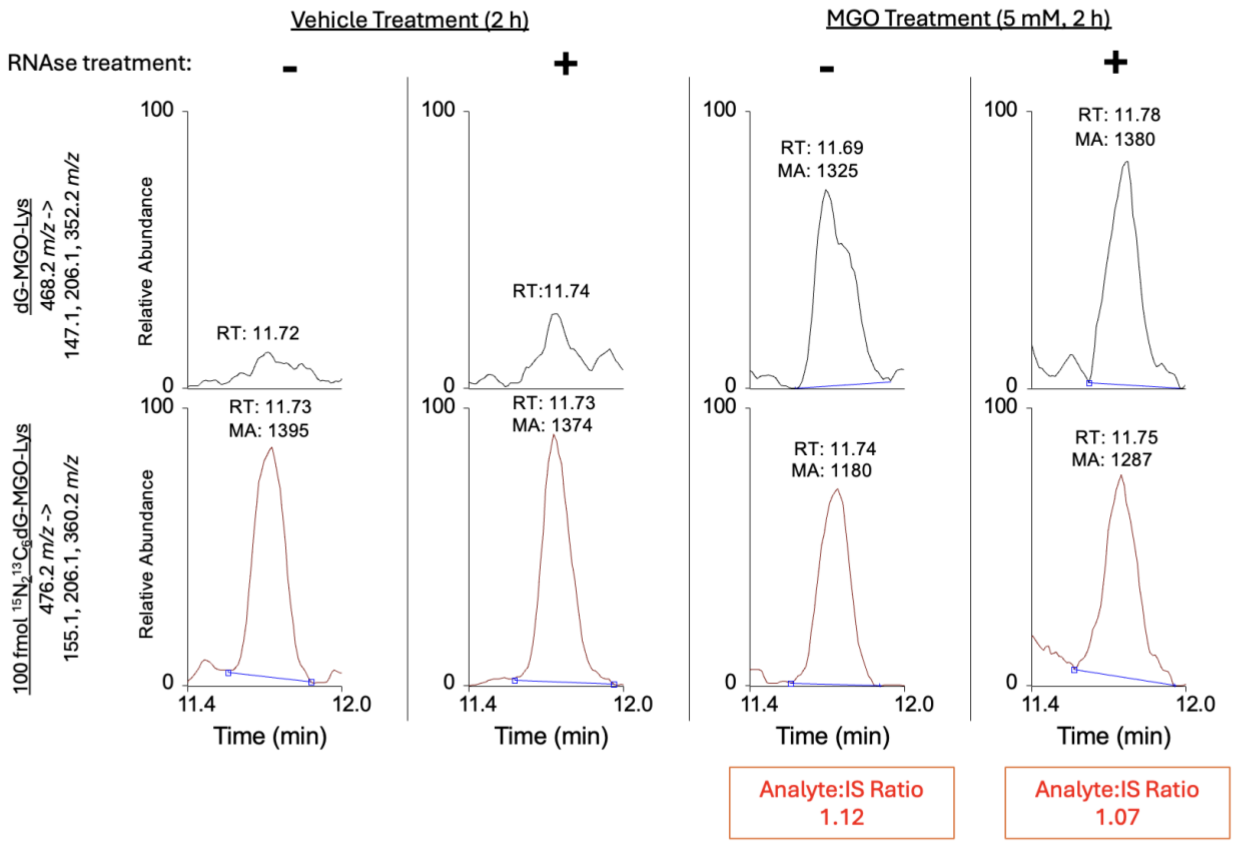


**Figure S5.** RNAse digestion effect on yield of dG-MGO-Lys adducts in HeLa cells. HeLa cells were treated with either vehicle or MGO (5 mM, 2 h) and 1 µg DNA samples were digested with RNAse (4 µL RNAse, Qiagen). Digested samples were precipitated before further pronase digestion. Representative extracted ion chromatograms from HPLC fraction B of unlabeled and isotope labeled dG-MGO-Lys standards are shown. Comparable analyte-to-IS ratios of 1.12 without RNAse treatment and 1.07 with RNase treatment are calculated from the extracted ion chromatograms.

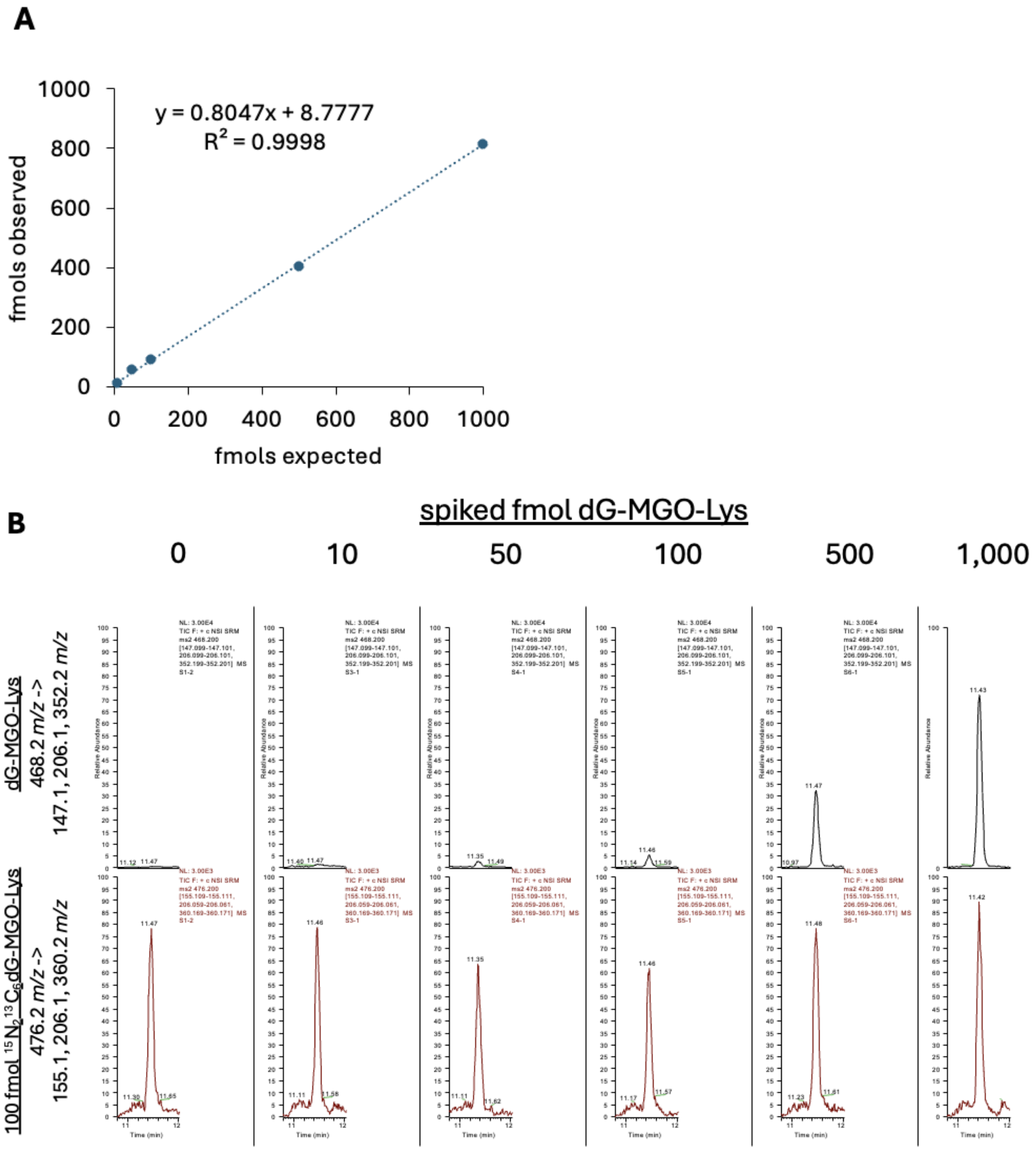


**Figure S6.** Calibration and method validation in DNA matrix**.** (**a**) Calibration curve for the authentic dG-MGO-Lys standards spiked in 1 µg unexposed DNA and processed according to the LC-MS/MS method. Observed signal (fmol) of unlabeled standard plotted against the signal for the isotope labeled standard across concentration range of 10-1000 fmol. Linearity, Limit of Detection (LOD), and Limit of Quantitation (LOQ) were calculated from these observations. (**b**) Extracted LC-MS/MS ion chromatograms for the unlabeled dG-MGO-Lys and the $^{15}N_2{}^{13}C_6$ –labeled internal standard at increasing dG-MGO-Lys concentrations.

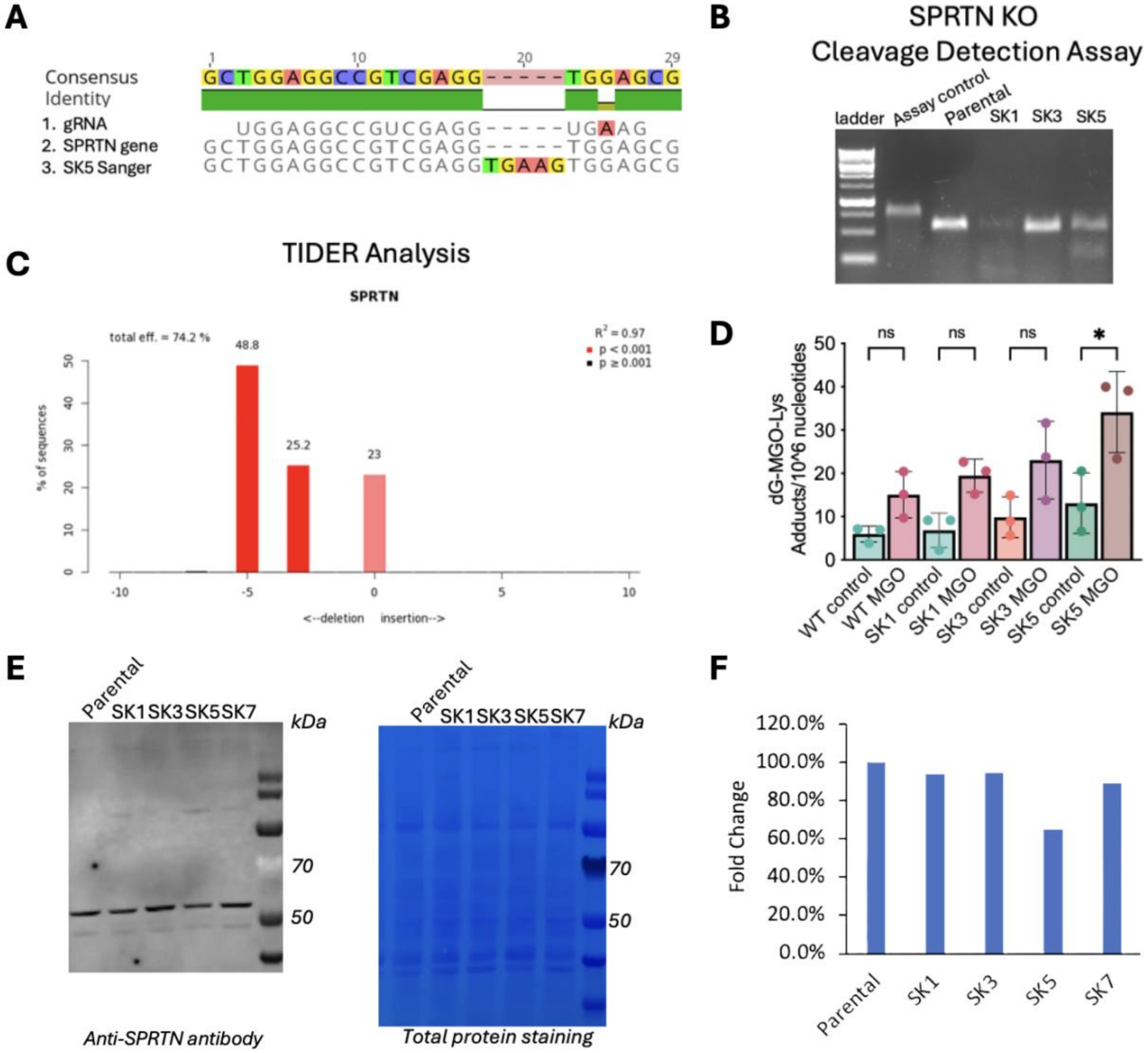


**Figure S7**. Validation of SPRTN knockout (KO) clones and their effects on dG-MGO-Lys adduct formation. (**a**) Sanger sequencing of targeted SPRTN locus showing alignment of CRISPR gRNA sequence, SPRTN gene consensus sequence, and SK5 clone Sanger sequencing results. (**b**) Cleavage detection assay confirming genomic editing in selected isolated clones (SK1, SK3, SK5). (**c**) TIDER analysis quantifying insertion/deletion events in SK5 clone, with a total editing efficiency of 74.2%. (**d**) LC–MS/MS quantification of dG-MGO-Lys adducts in wild-type (WT) and SPRTN KO cells under control and MGO treatment (5 mM, 2 h). n=3 (**e**) Western blot analysis using anti-SPRTN antibody and total protein staining verifying loss of SPRTN protein in KO clones. (**f**) Fold-change comparison of SPRTN protein expression in parental versus KO clones as observed by western blotting.

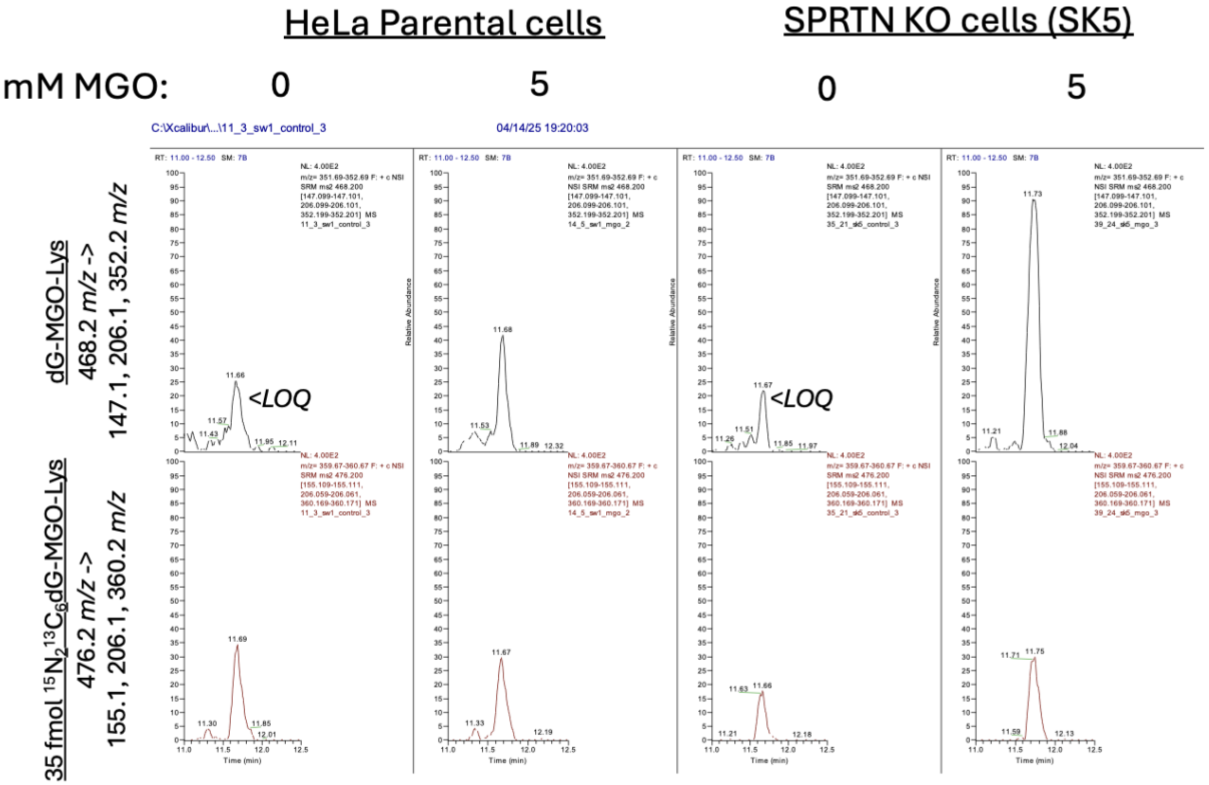


**Figure S8.** Representative LC–MS/MS ion chromatograms of dG-MGO-Lys adducts in HeLa parental and SPRTN KO (SK5) cells. Extracted ion chromatograms from HPLC fraction B of unlabeled dG-MGO-Lys (*m/z* 468.2 → 147.1, 206.1, 352.2) and isotope $^{15}N_2{}^{13}C_6$- dG-MGO-Lys standard (*m/z* 476.2 → 155.1, 206.1, 360.2) in parental and SPRTN KO cells exposed to 0 or 5 mM MGO for 2 h. dG-MGO-Lys signals were below the limit of quantitation (LOQ) in untreated samples.

### Materials and Methods

**Reagents and Materials.** Unless otherwise stated, all reagents and chemicals were used as received. Boc-protected isotope labeled $^{15}N_2{}^{13}C_6$ -L-lysine (98%), 2'-deoxyguanosine monohydrate (97%), magnesium chloride (99.9%), calcium chloride (97%), ammonium acetate (97%), sodium chloride (≥99%), dithiothreitol (DTT), ethanol (200 proof), acetonitrile (HPLC grade, 99.9%), isopropanol (HPLC grade, 99.5%), acetonitrile (LC-MS grade, 99.8%), water (LC-MS grade), formic acid (LC-MS grade, 99%), pronase, pierce™ BCA protein, and Quant-iT™ PicoGreen™ dsDNA assay kits were purchased from Thermo Fisher. Acetic acid (HPLC grade, 99.8%), carboxypeptidase Y, aminopeptidase M, alkaline phosphatase, and Nanosep 10k filters were obtained from Sigma-Aldrich. Phosphodiesterase I, phosphodiesterase II, and DNAse I were acquired from Worthington, while proteinase K was obtained from New England Biolabs. The unlabeled dG-MGO-Lys was prepared following our previously published procedure.[1] RNAse was obtained from Qiagen.

**STAR Assay.** The assay was adapted from the following reported literature method.[2] Cells were lysed in lysis buffer consisting of 50 mM Tris–HCl (pH 7.4), 1 mM EDTA, 150 mM NaCl, 1% Triton X-100, 0.5% deoxycholate (DOC), 0.1% sodium dodecyl sulfate (SDS), and protease inhibitor cocktail (Complete tablets, Easy Pack 04693116001, Roche) for 15 min on ice, followed by centrifugation at 16,000 x g for 15 min to remove the supernatant. The resulting pellet containing DNA and insoluble material was dissolved in buffer consisting of 6 M guanidinium–HCl, 10 mM Tris–HCl (pH 6.5), 20 mM EDTA, 4% Triton X-100, 0.1% SDS, and 1% dithiothreitol (DTT). DNA was precipitated with an equal volume of 200 proof ethanol. Pelleted DNA was recovered by centrifugation at 16,000 x g for 10 min, washed with 75% ethanol, and centrifuged again before discarding the supernatant and evaporating the residual ethanol. The dried pellets were resuspended in 1xTE buffer, and DNA was quantified using Quant-iT™ Picogreen, while protein content was assessed by BCA[3] or Bradford assays according to the manufacturer's instructions.[4]

**Western Blot**. Samples (30 µL) were mixed with 10 µL of 4 x loading buffer and adjusted to a final volume of 40 µL, followed by vortexing, boiling at 90 °C for 10 min, and centrifugation. Proteins were resolved on 4–12% SDS–PAGE gradient gels. Electrophoresis was carried out at 90 V for 1.5 h or until ladder separation was complete. For Coomassie staining, gels were washed three times with ultrapure water 5 min each, incubated in SimplyBlue stain (Thermo) for 1 h to overnight at room temperature. Gels were destained in ultrapure water until clear, with images acquired upon completion. For western blotting, proteins were transferred onto nitrocellulose membranes using the electrophoresis conditions of 20 V for 1 h. Membranes were blocked with 5% milk in TBST for 1 h at room temperature, and rinsed with TBST. Primary antibodies were added and membranes incubated overnight at 4 °C on a rotator. Following three TBST washes (5 min each), membranes were incubated with secondary antibodies at room temperature for 1 h. Membranes were washed again, and imaged using a LiCor instrument with ladder visualization at 700 nm.

**Cell Culture.** Cells were grown using Dulbecco's Modified Eagle's Medium (DMEM) containing 10% fetal bovine serum and 1% penicillin-streptomycin. Cells were maintained at 37 °C in a humidified incubator with 5% $CO_2$. GLO1 knockout HEK293 cells were received from Dr. James Galligan's laboratory. [5]

**Generation of SPRTN Knockout cells**. The TrueCut Cas9 Protein v2 was obtained from Thermofisher. The sgRNA and Cas9 nuclease was transfected into HeLa cells using Lipfectamine CRISPRMAX reagent in Opti-MEM I reduced serum medium (Thermo) according to the TrueCut Cas9 Protein v2 manufacturer's instructions. The sgRNA sequence (CACGCTCCACTTCACCTCGACGG) targeted the first exon of SPRTN. This gRNA was previously used to generate a SPRTN knockout cell line.[6] 48 h post transfection, cells were seeded in a 96-well plate (one cell/well). Cells were then grown to confluency. After single cell clones were expanded into 6 well plates, 3 distinct CRISPR-targeted HeLa cell clones (each grown from a single cell) were tested for CRISPR cleavage activity. We used the GeneArt Genomic Cleavage Detection Kit (Thermo), Sanger sequencing, and western blotting for SPRTN protein expression (antibody: 144-62088-20, Ray Biotech). A control treated clone was used as a negative control. Primers for the cleavage detection kit and follow-up Sanger sequencing included Primer 1 Forward: CTGTGATCCTGGCAACGATG, Primer 1 Reverse: TCCAAAACCAGACTCCCCTC, Primer 2 Forward: TGACTTGATGTTGGCACTGC, Primer 2 Reverse: GAAGACAGAAACACGCCGAG. Sanger sequencing results were processed with TIDE software (version 5.0.2). [7]Approximately 50% of the sequences exhibited a five-nucleotide addition in exon 1 of SPRTN, resulting in a premature stop codon. HeLa cells, a cancer cell line, has multiple copies of chromosome 1, the location of the SPRTN gene. These results suggest that the SPRTN gene is only partially knocked-out, consistent with the partial loss of SPRTN protein abundance observed by western blotting.

**Synthesis of labeled dG-MGO-Lys Internal Standard.** The reaction was carried out as previously described in the literature.[8] To a stirring solution of dG (5 mM) in PBS (pH 7.4), boc protected isotope-labeled $^{15}N_2{}^{13}C_6$ - L-Lys (5 mM), and MGO (5 mM) were added and the reaction mixture was incubated at 37 °C for 3 days. The reaction was monitored via HPLC where 100 µL injections of the reaction mixture were separated on a Synergi 4 µm Hydro-RP 80 Å, LC Column 250 x 4.6 mm with buffer A: $H_2O$ and buffer B: ACN over the following gradient: 1 to 10%B over 15 min, then 10 to 20% B over 5 min, a hold at 20% B for 5 min, then 20 to 95% B over 10 mins, and a 5 min hold at 95% B. Throughout the run, 1 min fractions were collected and were further analyzed on a Thermo Scientific LTQ XL linear ion trap mass spectrometer to detect labeled dG-MGO-Lys crosslinked product. Fractions containing labeled dG-MGO-Lys were then subjected to the hydrogenation deprotection reaction. The final product was isolated via HPLC on an analytical column, Synergi 4 µM Fusion-RP 80 250 x 4.6 mm (part number: 00g-4375-e0) at a flowrate of 1 mL/min with the same gradient as above.
